# The infectious salmon anaemia virus esterase prunes erythrocyte surfaces in infected Atlantic salmon and exposes terminal sialic acids to lectin recognition

**DOI:** 10.1101/2023.01.25.525626

**Authors:** Johanna Hol Fosse, Adriana Magalhaes Santos Andresen, Frieda Betty Ploss, Simon Chioma Weli, Inger Heffernan, Subash Sapotka, Krister Lundgård, Raoul Valentin Kuiper, Anita Solhaug, Knut Falk

## Abstract

Many sialic acid-binding viruses express a receptor-destroying enzyme (RDE) that removes the virus-targeted receptor and limits viral interactions with the host cell surface. Despite a growing appreciation of how the viral RDE promotes viral fitness, little is known about its direct effects on the host. Infectious salmon anaemia virus (ISAV) attaches to 4-*O*-acetylated sialic acids on the surface of Atlantic salmon epithelial, endothelial, and red blood cells. ISAV receptor binding and destruction are effectuated by the same surface molecule, the haemagglutinin esterase (HE). We recently discovered a global loss of vascular 4-*O*-acetylated sialic acids in ISAV-infected fish. The loss correlated with the expression of viral proteins, giving rise to the hypothesis that it was mediated by the HE.

Here, we report that the capacity to bind new ISAV particles is also progressively lost from circulating erythrocytes in infected fish. Furthermore, salmon erythrocytes exposed to ISAV *ex vivo* lost their capacity to bind new ISAV particles. The loss of ISAV binding was not associated with receptor saturation. Moreover, upon loss of the ISAV receptor, erythrocyte surfaces became more available to the lectin wheat germ agglutinin, suggesting a potential to alter interactions with endogenous lectins of similar specificity. The pruning of erythrocyte surfaces was inhibited by an antibody that prevented ISAV attachment. Furthermore, recombinant HE, but not an esterase-silenced mutant, was sufficient to induce the observed surface modulation. Our results directly link the ISAV-induced erythrocyte modulation to the hydrolytic activity of the HE and show that the observed effects are not mediated by endogenous esterases.

Our findings are the first to directly link a viral esterase to extensive host cell surface modulation in infected individuals. This raises the question of how common the phenomenon is among sialic acid-binding viruses. It is also relevant to ask if the altered sialic acid landscape of the affected cells influences host biological functions with relevance to viral disease.

## Introduction

Sialic acids are highly diverse (>80 derivatives known to date) and typically present on the outermost ends of glycan chains attached to plasma membrane-anchored proteins or lipids [1]. Many sialic acid-binding viruses express a receptor-destroying enzyme (RDE) that removes the virus-targeted receptor and limits viral host cell attachment [2]. An appropriate balance between viral receptor-binding and receptor-destroying activities promotes viral fitness. The RDE supports both early and late steps in the infectious cycle: First, RDE activity destroys decoy receptors in mucus and reduces the density of virus-targeted sialic acids on cell surfaces, and this appears to help virus particles reach sites on the plasma membrane that favour viral entry [3–6]. Second, upon budding from the plasma membrane, RDE activity is required to prevent aggregation of viral particles and allow the release of new infective virus [7]. The importance of viral receptor destruction can be exemplified by the strong reduction of influenza virus replication by compounds that inhibit its RDE, the neuraminidase [6]. In addition, attachment interference resulting from viral receptor destruction mediates host cell resistance to superinfection [8–11].

Despite the growing appreciation of the role of the viral RDE in the infectious cycle, little is known about its direct effects on the host. First, how extensive is the RDE-mediated modulation of target cell surfaces in an infected individual? Second, considering that cell surface sialic acids modulate a range of cellular functions, including the activation of immune responses [12–14], could the loss of sialic acid viral receptors influence host biological functions with relevance to viral disease?

Infectious salmon anaemia virus (*Isavirus salaris*, ISAV) is an enveloped, segmented, single-stranded, negative polarity RNA virus of the *Orthomyxoviridae* family. ISAV contains eight genomic segments that encode at least 10 proteins [15–17]. Amongst these is the dual-function surface glycoprotein haemagglutinin esterase (HE) that is responsible for both binding to and hydrolysis of the 4-*O*-acetylated sialic acid identified as the ISAV receptor [18, 19]. Infection with pathogenic ISAV causes disease (infectious salmon anaemia, ISA) in farmed Atlantic salmon (*Salmo salar* L.) [20], and has led to vast economic losses in all major salmon-producing countries.

Vascular endothelial cells are the main target cells of pathogenic ISAV and support the generation of new virus particles that are released into the blood stream [21, 22]. Moreover, clinical signs of ISA are compatible with a viral sepsis-like breakdown of central vascular functions, including petechial bleeds, vascular leakage, and focal necrosis [20]. We recently revealed a global loss of vascular 4-*O*-acetylated sialic acids in ISAV-infected fish that correlated with the expression of viral proteins [23]. While our findings suggested that the esterase activity of the HE could be involved in the modulation of host cell surfaces, we could not completely exclude the involvement of endogenous host esterases.

Several lines of reasoning made us curious if erythrocyte surfaces were modulated in a similar manner in infected fish: First, nucleated fish erythrocytes are targeted by ISAV, but show limited, if any, permissiveness to infection [24]. Consequently, any surface modulation would be independent of the cellular expression of viral proteins. Second, prior to the onset of sepsis-like clinical signs and mortality, ISAV-infected fish typically develop anaemia [24, 25]. In general, sialic acids and their 9-*O*-acetylation are involved in regulating the circulating half-life of erythroid-lineage cells in other species [26–29]; hence, a modulation of erythrocyte sialic acids could potentially contribute to the pathogenesis of ISA. Finally, erythrocytes are accessible to sampling and *ex vivo* manipulation, facilitating exploration of the underlying mechanisms.

We found that erythrocytes in infected fish, similar to endothelial cells, progressively lose the ability to bind new ISAV particles. By exposing erythrocytes from non-infected fish to ISAV and recombinant proteins *ex vivo*, we further revealed that the loss of ISAV binding was not due to saturation of the receptor, but associated with pruning of surface sialic acids by the ISAV esterase. These findings expand on our recent study that showed loss of vascular 4-*O*-acetylated sialic acids in infected fish. First, by providing direct mechanistic evidence that the surface modulation is caused by ISAV esterase activity, rather than endogenous host esterases. Second, by demonstrating that cells can be extensively modulated by RDE activity, despite not permitting viral replication. Our observations raise the questions of whether RDEs of other sialic acid-binding viruses modulate target cell populations to the same extent, and if such surface modulations influence biological processes in infected hosts.

## Results

### Disease and viraemia in experimentally infected fish

Atlantic salmon (n=47) were challenged by immersion for two hours with ISAV (Glesvaer/2/90, 10^3.75^ TCID_50_/mL) and subsequently maintained in a fresh water flow-through system at 12 °C for the duration of the trial (Fig 1A). The first mortality in the infected fish group occurred 15 days post infection (dpi), with mortality rapidly increasing over the next days (Fig 1B). The trial was terminated 18 dpi, upon reaching its pre-determined end point of 40% cumulative mortality. No fish in the non-infected group (n=26) died during the trial. Infected fish developed anaemia 12 dpi (Fig 1C), prior to other clinical signs. Histological examination of haematoxylin & eosin-stained sections of formalin-fixed paraffin-embedded tissues revealed increased erythrophagocytosis in head kidney and spleen of infected fish that coincided with the onset of anaemia (S1A-C Fig), suggesting that erythrocytes were removed from the circulation at an increased rate. Viraemia, measured by ISAV segment 8 RNA and infective particles in the blood, was detected in 3 out of 4 fish at the earliest sampling point (4 dpi) and all infected fish at subsequent samplings. Blood viral loads peaked and plateaued 14 dpi (Fig 1D), with no significant decline within the trial period (S1 Table). Levels of viral RNA in the head kidney peaked and plateaued earlier, 10 dpi (S1C Fig), in line with the assumption that circulating ISAV particles originate from extensive replication in vascular endothelial cells [21, 22].

**Fig 1:**
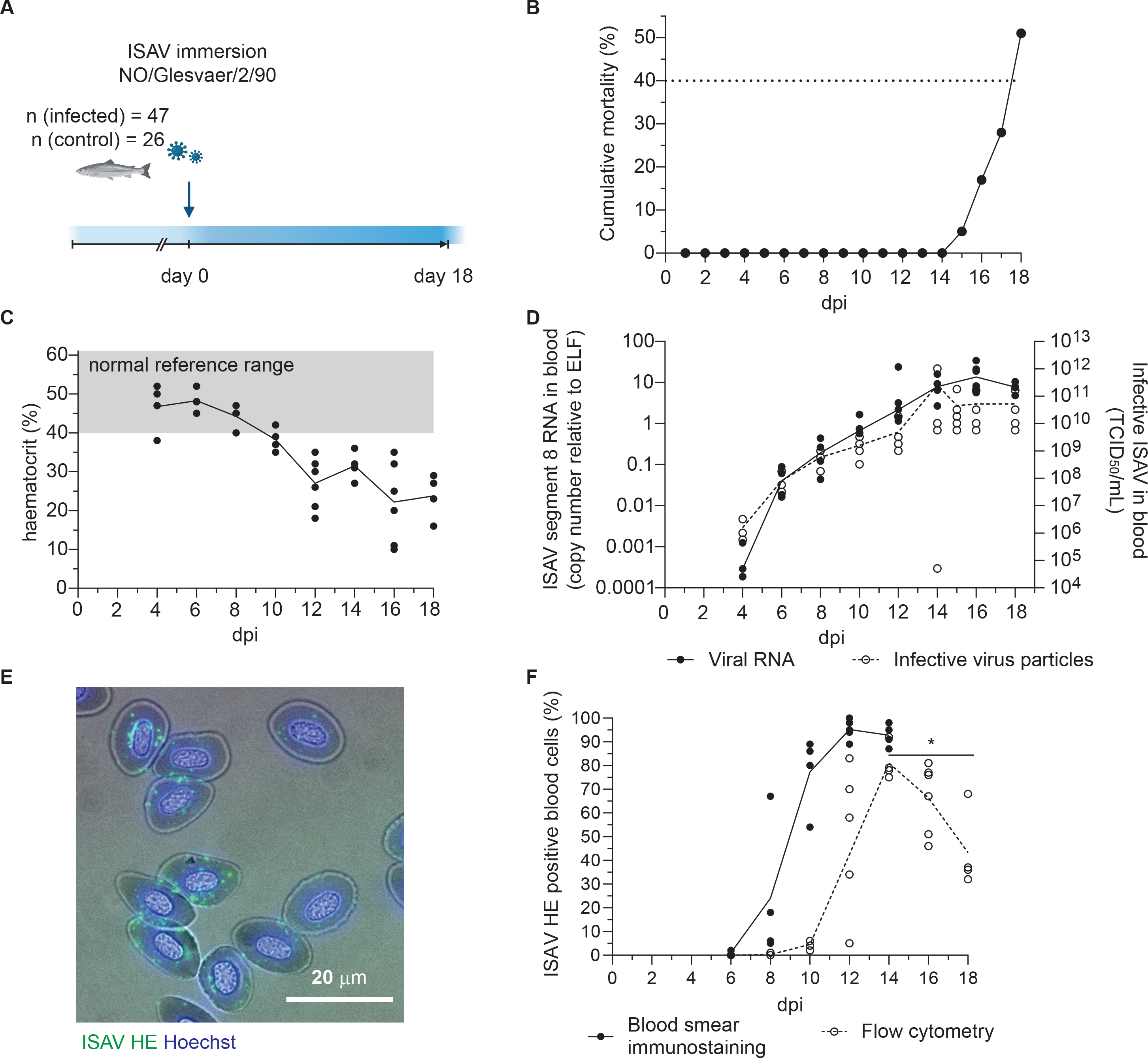
Disease and viraemia in experimentally infected fish. (A) Outline of the infection trial. Atlantic salmon were challenged with ISAV (NO/Glesvaer/2/90, 2 hours immersion) and followed for 18 days. The illustration was created in BioRender.com. (B) Cumulative mortality in the infected fish group. No deaths occurred in the non-infected group. The stippled line indicates the prior determined humane end point. (C) Haematocrits (HCT) of infected fish. Data points represent individual fish, and the line connects median values. The reference range (grey shading) is based on blood samples from 25 non-infected fish (mean +/− 2 standard deviation). (D) Viral RNA and infective particles in blood were measured by qPCR targeting ISAV segment 8 (left y-axis, black dots and line) and titration on ASK cells (right y-axis, open circles, stippled line). Data points represent individual fish, lines connect median values. (E) Representative micrograph of an acetone-fixed blood smear from an infected fish (12 dpi) immunostained for HE (clone 3H6F8, green). Nuclei are counterstained with Hoechst 33342 (blue), and bright field contrast shows the outline of the cells. The scale bar measures 20 μm. (F) The percentage of HE-positive cells in acetone-fixed blood smears were counted manually (black dots and line). Non-permeabilised PFA-fixed blood cells from infected fish were immunostained for HE, and the percentages of positive cells were determined by flow cytometry (open circles, stippled line). Data points represent individual fish. Lines connects median values. *p<0.05: flow cytometry data from 14, 16, and 18 dpi were compared by Kruskall-Wallis test and Dunn’s multiple comparison (S1 Table).

The attachment of ISAV to erythrocyte surfaces was first evaluated by manual counting of immunostained blood smears, detecting ISAV-positive erythrocytes in 2 out of 4 fish 6 dpi and all fish at subsequent samplings (Fig 1E-F, black dots). Blood smears obtained at 16 and 18 dpi were not of sufficient technical quality for analysis. Flow cytometry provided a less sensitive, but more objective method for quantifying the percentage of erythrocytes with surface-bound ISAV, and was performed at all time-points (Fig 1F, open circles, stippled line). By flow cytometry, ISAV-positive cells were detected in 1 out of 4 fish 8 dpi and all fish at subsequent samplings, with the fraction of positive cells peaking 14 dpi. A decline in ISAV-positive cells was detected from 14 to 18 dpi (S1 Table). The percentage of ISAV-positive cells assessed by either flow cytometry or immunostaining of blood smears correlated with infective titres (Spearman r = 0.8353 and 0.8302; p<0.0001 and p<0.001, respectively).

### The ISAV receptor is lost from circulating erythrocytes in infected fish

Next, we assessed the distribution of the ISAV receptor in tissue sections and membrane-enriched fractions of erythrocytes from infected fish. We performed a virus binding assay where ISAV antigen produced in ISAV-infected cells was used as the primary probe [21] (Figure 2A). Consistent with our recent observations [23], heart vascular endothelial cells lost their capacity to bind new ISAV particles 10 dpi and did not recover (Fig 2B-C). Despite ISAV remaining attached to circulating erythrocytes throughout the course of infection (Fig 1F), a similar loss of capacity to bind new ISAV particles was observed 10 dpi onwards (Fig 2D-F). New ISAV binding to heart sections and erythrocyte membranes correlated in individual fish (Spearman’s correlation coefficient = 0.7494, p<0.0001, Table S1). These findings reveal that erythrocytes in ISAV-infected fish lose the ability to bind new ISAV particles as the infection progresses in a similar manner to vascular endothelial cells [23], suggesting loss of the host cell surface receptor.

**Fig 2:**
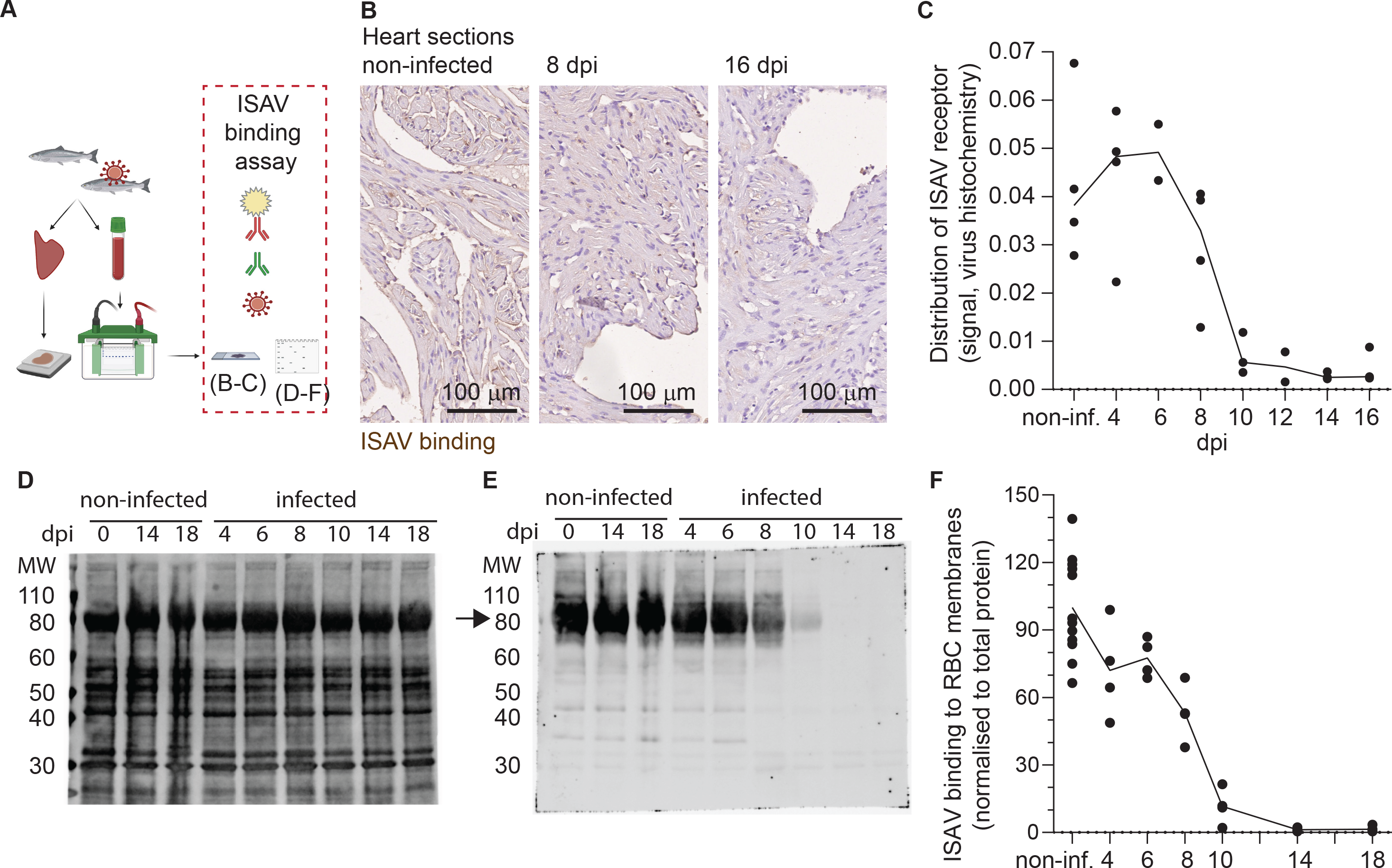
Circulating erythrocytes in infected fish lose the ability to bind new ISAV. (A) The distribution of the ISAV receptor in heart tissues and membrane-enriched erythrocyte fractions was evaluated by serial incubation with ISAV antigen, mouse IgG_1_ targeting HE (clone 3H6F8), fluorescence- or HRP-conjugated secondary antibodies, and substrate if relevant. (B) Representative micrographs showing the ISAV receptor (brown) in hearts of non-infected and infected fish (8 and 16 dpi). Sections are counterstained with haematoxylin. Scale bars measure 100 μm. (C) Quantification of ISAV receptor signal in scanned heart sections. (D-E) Membrane-enriched erythrocyte lysates were separated by SDS-PAGE and blotted to nitrocellulose membranes, before evaluation of ISAV binding as described above. Representative blots show (D) total protein and (E) ISAV binding (arrow) to samples from non-infected fish harvested 0, 14, and 18 dpi and infected fish harvested 4, 6, 8, 10, 14, and 18 dpi. (F) Quantification of signal (4 fish per time point) normalised to total protein. (C and F) Data points represent individual fish. Lines connect median values.

### The erythrocyte loss of ISAV binding is not mediated by saturation of the ISAV receptor

To understand the mechanisms behind the observed ISAV receptor loss in infected fish, we next exposed density-purified erythrocytes from non-infected fish to supernatants from ISAV-infected cells. After identifying a virus dose that did not saturate the ISAV binding capacity (10^5^ TCID_50_ per 2×10^7^ cells, Fig 3A), we evaluated the ISAV binding capacity of cells exposed to this non-saturating dose for 20 hours. Reflecting the situation in infected fish, we found that prior exposure to ISAV abolished subsequent ISAV binding to the membrane-enriched fractions (Fig 3B-C).

**Fig 3:**
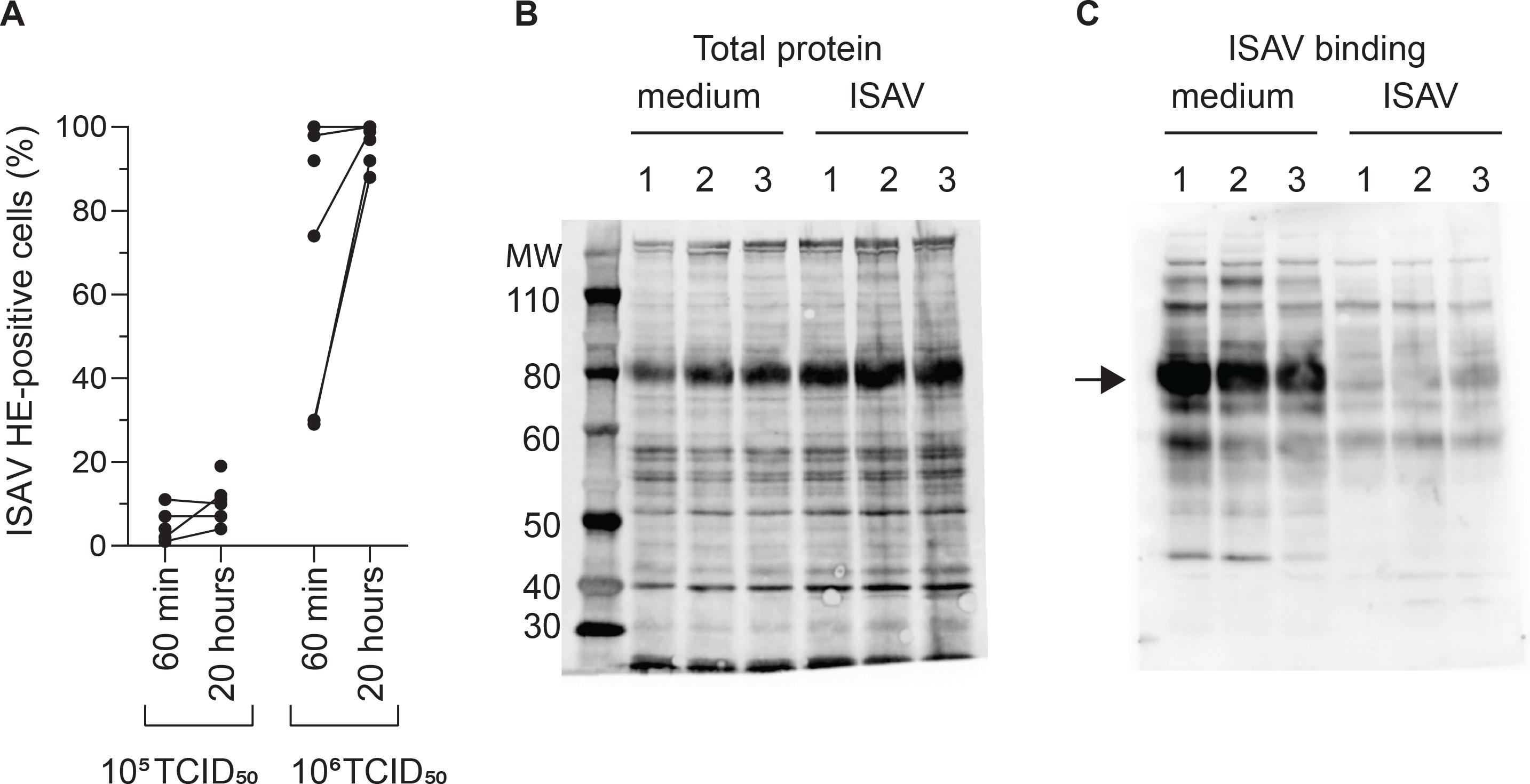
Loss of ISAV binding is not mediated by saturation of its cellular receptor. (A) Density-purified erythrocytes isolated from non-infected fish were incubated with ISAV at the indicated dose (per 20 million cells) and given duration, and the percentage of HE-positive cells was quantified by flow cytometry. Data points show measurements in cells from individual fish, with lines connecting values from the same fish when relevant. (B-C) Membrane-enriched erythrocyte lysates from three fish (1, 2, 3) exposed to medium or ISAV (10^5^ TCID_50_ per 20 million cells, 20 hours) were separated by SDS-PAGE and blotted to nitrocellulose membranes. (B) Blots were stained for total protein by Revert700 Total Protein Stain, (C) and the level of ISAV receptor was evaluated by serial incubation with ISAV antigen, mouse IgG_1_ targeting HE (clone 3H6F8), HRP-conjugated secondary antibody, and substrate. The assay was repeated twice with identical results. The arrow points to the band representing ISAV binding in medium-treated erythrocytes.

This illustrates that exposure to ISAV-containing supernatants is sufficient to reproduce the loss of ISAV binding observed in infected fish, and that the loss of binding is not due to receptor saturation.

### The loss of ISAV binding is accompanied by increased availability to sialic acid-binding lectins

Loss of 9-*O*-acetylation can make sialic acids more available to endogenous sialic acid-binding immunoglobulin-like lectins (siglecs), probably by reducing steric hindrance [29–31]. Similarly, 4-*O*-acetylation prevents binding of the plant lectin WGA to a range of α2,3-linked sialic acids [32]. Four members of the siglec family (siglec 1, 2, 4, and 15) are conserved in all vertebrates, including Atlantic salmon [33]. As the binding specificities of Atlantic salmon siglecs have not been characterised, we here used WGA as a proxy for testing if the loss of the ISAV receptor could modulate interactions between lectins and Atlantic salmon erythrocytes.

We found that exposure to both low (10^5^ TCID_50_ per 2×10^7^ cells) and high (10^6^ TCID_50_ per 2×10^7^ cells) doses of ISAV increased the binding of WGA to erythrocytes (Fig 4A). The effect was evident as soon as 60 min after exposure. At this time, the low dose effect was less prominent, but after 20 hours, the low and high doses of virus increased WGA binding to a similar level (Fig 4A, S1 Table). Only the high dose of ISAV increased binding of the Sambucus nigra lectin (SNA), which specifically targets α2,6-linked sialic acids [32] (Fig 4B).

**Fig 4:**
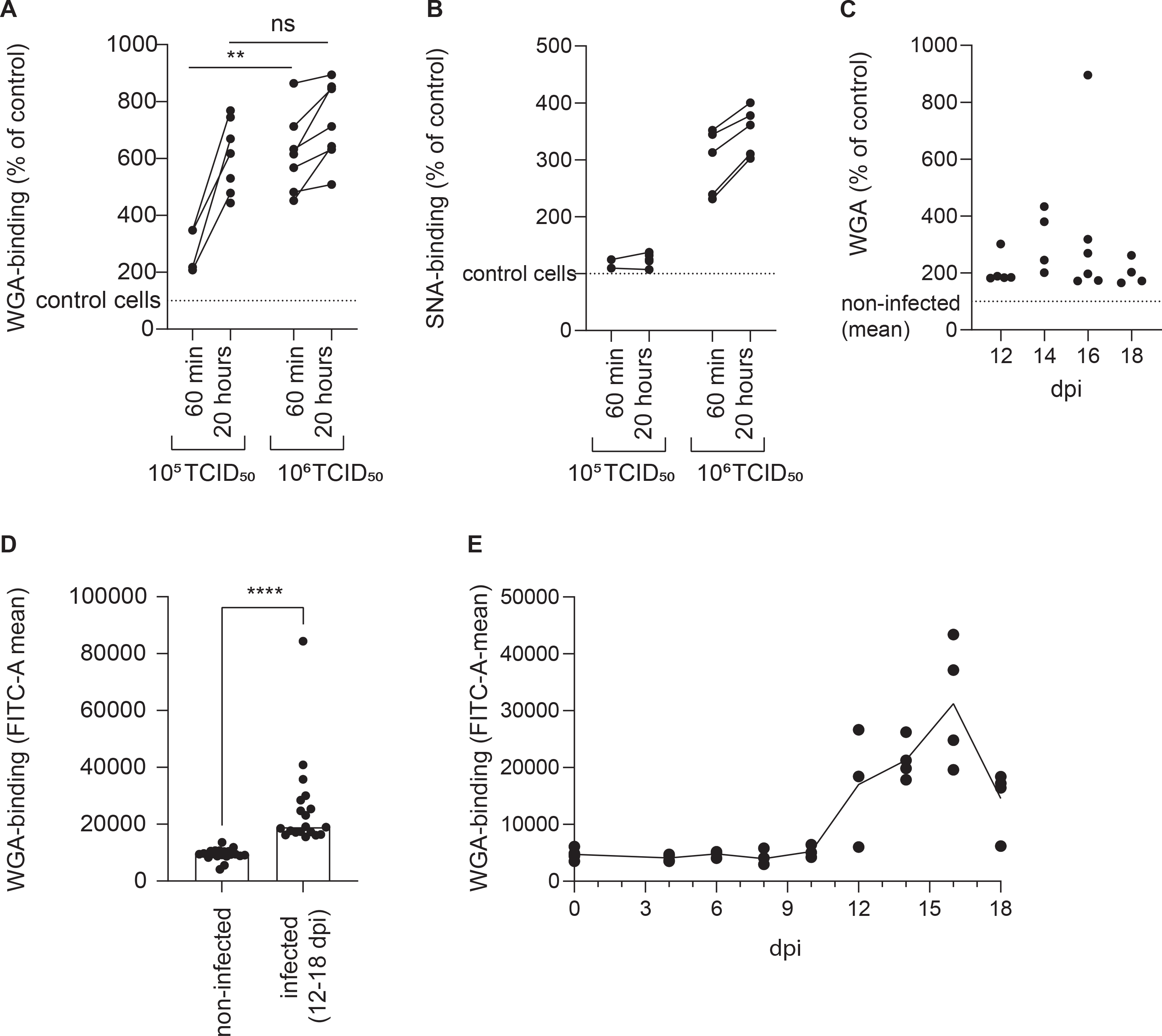
The loss of ISAV binding is accompanied by increased availability to sialic acid-binding lectins. The lectin-binding capacity of erythrocytes was evaluated by flow cytometry. (A-B) Density-purified erythrocytes isolated from non-infected fish were incubated with ISAV at the indicated dose (per 20 million cells) and given duration, and WGA-binding (A) and SNA-binding (B) was quantified in live cells by flow cytometry. Data points show measurements in cells from individual fish, with lines connecting values from the same fish. ** p<0.01, Mann-Whitney U. (C-D) WGA-binding to PFA-fixed blood cells from infected fish was quantified by flow cytometry. Data points show values in individual fish. **** p<0.0001, Mann-Whitney U. (E) Density-purified erythrocytes were incubated with plasma from infected fish (10 μL/10^6^ cells) and incubated overnight before quantification of WGA-binding by flow cytometry. Data points show cells incubated with plasma from individual fish (4 per time point). The line connects medians.

Erythrocytes of infected fish (12-18 dpi) also bound WGA more efficiently than cells from non-infected individuals harvested at the same time (Fig 4C-D), confirming the *in vivo* relevance of our finding. Finally, incubation of erythrocytes from healthy fish with plasma from infected fish increased their binding to WGA (Fig 4E). Due to limited materials being available, SNA-binding was not tested in samples from infected fish.

### The modulation of erythrocyte surfaces is mediated by the ISAV esterase

A monoclonal antibody (mouse IgG_1_ clone 9G1F10A) that inhibited cellular ISAV attachment and haemagglutination (S2 Fig) prevented the increase in WGA and SNA binding (Fig 5A-C). Similarly, the increase in WGA-binding observed when erythrocytes were incubated with plasma from infected fish, was strongly reduced when the plasma samples were pre-incubated with the neutralising antibody (Fig 5D). Finally, incubating erythrocytes with recombinant HE showed that this protein was sufficient to induce an increase in WGA-binding (Fig 5E-F). However, when its esterase activity was silenced by alanine mutation of the catalytic serine (S32) [34], the HE-mediated increase in WGA was abolished (Fig 5E-F).

**Fig 5:**
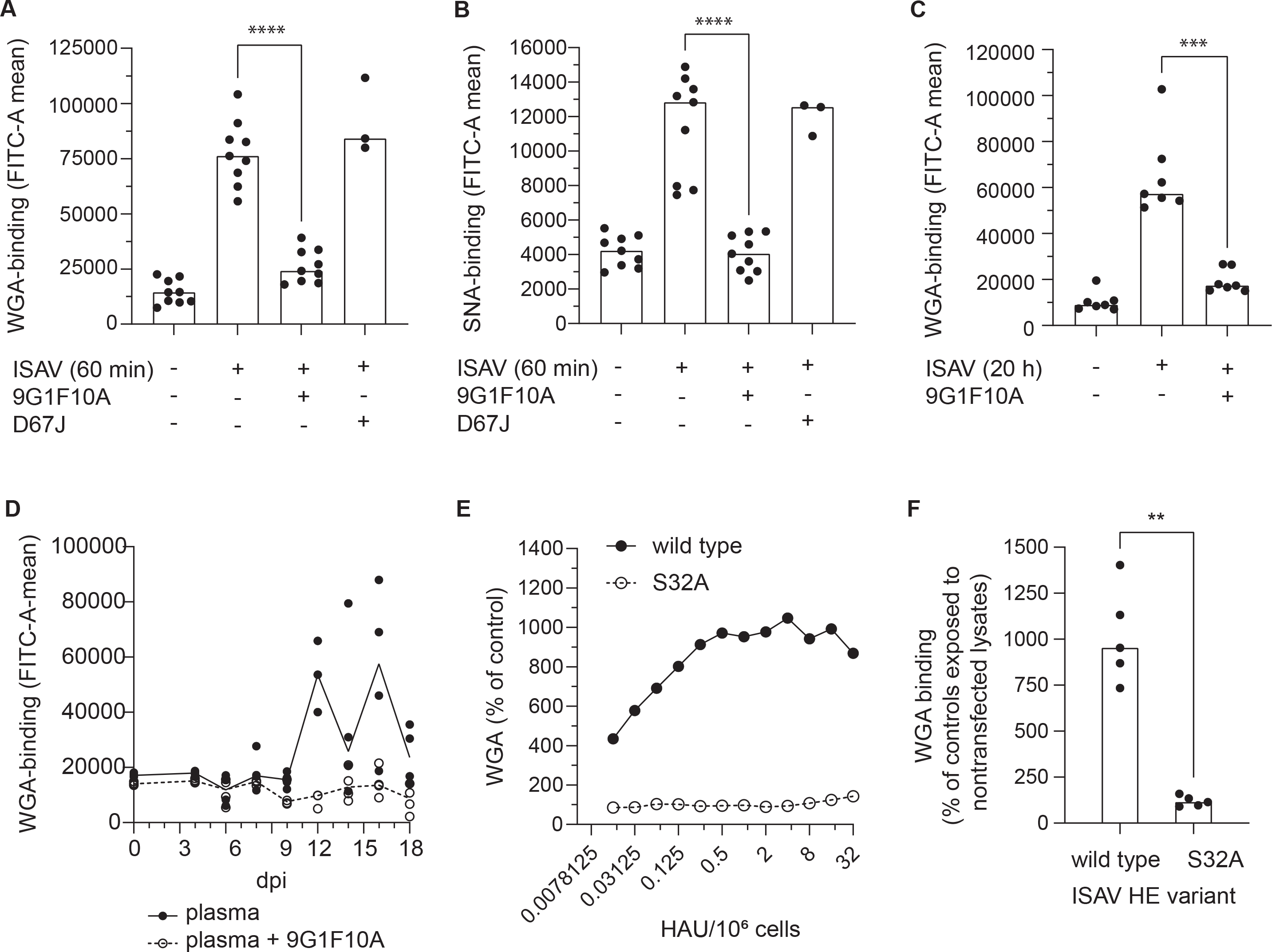
The pruning of erythrocyte surface is mediated by the ISAV esterase. (A-C) Virus supernatants were pre-incubated with antibodies targeting HE (9G1F10A) or the nucleoprotein of influenza A virus (D67J) (both: 5 μg/mL, 30 min, RT) before inoculation with erythrocytes (A-B: 10^6^ TCID_50_ per 20 million cells, 60 min; C: 10^5^ TCID_50_ per 20 million cells, 20 hours). (A, C) WGA-binding and (B) SNA-binding, measured by flow cytometry. (D-E) WGA-binding to erythrocytes exposed to wild type or esterase-silenced (S32A) HE at 1 HAU per 10^6^ cells for 20 hours. (A-D and F) Each data point represents blood cells isolated from one fish. Bars and line show median values. **p<0.01, ***p<0.001, ****p<0.0001, Mann Whitney U.

Together, these findings suggest that sialic acids on ISAV-exposed erythrocytes become more available to lectin binding; that attachment of ISAV to the cellular surface is required for this effect to take place; and that the ISAV-induced surface modulation is at least in part mediated by the hydrolytic activity of the viral RDE.

Our collected results support the conclusion that the ISAV RDE extensively removes 4-*O*-acetylated sialic acids from target cell surfaces in infected hosts and suggest a potential to influence interactions with endogenous lectins.

## Discussion

We recently discovered that the ISAV receptor disappears from the vascular surface of ISAV-infected Atlantic salmon [23]. Here, we confirm and expand on that study, by showing that circulating erythrocytes in infected fish also lose the ability to bind new ISAV particles and identifying a key role for the viral esterase in this process. Our findings support a model where the HE first binds 4-*O*-acetylated sialic acids on the cell surface to allow ISAV attachment, and then causes extensive hydrolysis of adjacent non-bound 4-*O*-acetylated sialic acids. The loss of cell surface 4-*O*-acetylated sialic acids limits further ISAV binding and makes cell surface sialic acids more available to lectins, but does not affect already-bound ISAV particles.

Despite being nucleated, and in contrast to endothelial cells [21], only a minor fraction of Atlantic salmon erythrocytes supports ISAV protein production [24]. Our findings therefore illustrate that the loss of viral receptor does not depend on viral replication and suggest that the cell surface is modulated at an early stage of the infectious cycle.

The loss of erythrocyte ISAV binding could be attributed to the hydrolytic activity of HE: First, the effect was not associated with receptor saturation; suggesting that it was mediated by hydrolysis of 4-*O*-acetyl sialic acids, like in endothelial cells [23]. Second, the loss of ISAV receptor from vascular endothelial cells and erythrocytes correlated in individual fish, indicating a common denominator. Third, exposing normal erythrocytes to plasma from infected fish reproduced the surface modulation, suggesting that this denominator was present in the blood. The plasma-induced surface modulation was inhibited by an antibody that prevented ISAV attachment, strengthening the assumption that circulating ISAV particles were responsible for the effect, rather than secreted host esterases. ISAV-dependent surface modulation was also observed when erythrocytes from non-infected fish were exposed to supernatants from infected cells. Finally, exposure to recombinant HE was sufficient to induce the same surface modulation in normal erythrocytes, but only when the HE hydrolytic activity was preserved.

Even though erythrocytes from infected fish no longer supported the attachment of new virus particles, they retained surface-bound ISAV throughout the trial, with only a small decline over the last four days. Several studies have illustrated that the dissociation of ISAV from Atlantic salmon erythrocytes is limited. The isolate used in the current study, Glesvaer/2/90, was the first ISAV isolate propagated [35] and has since been used in many of the key publications that characterise ISAV proteins, including HE [16–19, 34]. The isolate efficiently hydrolyses both free and glycosidically bound 4-*O*-acetylated sialic acids [18], and both horse, rabbit, rainbow trout, cod, and crucian carp erythrocytes elute in the expected manner after Glesvaer/2/90-mediated agglutination [16, 17]. Yet, the virus does not permit elution of agglutinated Atlantic salmon erythrocytes [16, 17]. Interestingly, the same lack of elution is seen when Atlantic salmon erythrocytes are agglutinated with influenza C virus [17], which binds and hydrolyses 9-*O*-acetylated sialic acids [36, 37]. This suggests that the lack of virus dissociation might be due to specific properties of Atlantic salmon erythrocytes.

The discrepancy between hydrolytic activity and virus dissociation indicates that the release of ISAV from the cell surface depends more on the on-off rate of the initial receptor engagement than the viral esterase hydrolytic activity. In other words, the ISAV esterase appears to limit the initiation of new attachments, rather than inducing the dissociation of already-bound ISAV particles. Nevertheless, the hydrolysis of adjacent receptors will reduce the local receptor density around the site initially bound by the virus, limiting the possibility to reattach to the cellular surface after release of the initial attachment [3, 4].

The erythrocyte surface modulation in infected fish that abrogated further ISAV attachment also increased the binding of the plant lectin WGA. The hydroxyl-group of sialic acid carbon 4 contributes to determining the specificity of WGA [38]. Also, the binding of WGA to a range of α2,3-linked sialic acids is inhibited by 4-*O*-acetylation [32]. It is therefore not surprising that the ability to bind WGA increases when the ISAV receptor is lost. The effect on SNA-binding was more difficult to predict. SNA is highly specific to α2,6-linked sialic acids, and its binding is not inhibited by 9-*O*-acetylation [32]. While we have not identified any previous studies of how 4-*O*-acetylation influences SNA-binding, our findings suggest that this modulation could limit interactions between SNA and its ligand on Atlantic salmon erythrocytes, presumably α2,6-linked sialic acids.

In addition to increasing the knowledge of how ISAV interact with target cell surfaces in the early phase of the infectious cycle, our observations raise two important questions:

*First, does the RDE-induced cell surface-modulation influence biological functions in the host?* In viral infection, it is essential for a successful outcome that the immune response is powerful enough to eliminate the viral infection, but controlled and specific enough to avoid unsustainable collateral damage to the host. In addition to being exploited by numerous microbes for attachment [2], sialic acids regulate host tolerance and immune activation by interactions with siglec-expressing immune cells [12–14, 39]. These interactions can be modulated by sialic acid *O*-acetylation [40–42]. Sialic acid de-acetylation, which we have identified on erythrocyte and vascular surfaces in ISAV infected fish, typically enhances siglec interactions [40], and the biological end-result will depend on the specific siglec and cell type involved. Atlantic salmon siglecs and other sialic acid-binding molecules are poorly characterised. Hence, it is difficult to predict the effects of the ISAV RDE-induced loss of vascular and erythrocyte 4-*O*-acetylated sialic acids. However, in the light of the pathogenesis of ISA, this subject deserves further investigation. ISA is characterised by a severe reduction in the number of circulating erythrocytes, circulatory collapse, and surprisingly low levels of perivascular inflammation, despite a strong vascular endothelial expression of viral proteins [20, 21], Amongst several interesting observations that may be relevant to ISA pathogenesis is that loss of 9-*O*-acetylation enhanced interactions between murine erythroid leukaemia cells and siglec-1, promoting erythroid cell retention in the spleen and liver [29]. Furthermore, loss of *O*-acetylated sialic acids promoted the activation of natural killer cell inhibitory siglecs and protected human cancer cell lines from cytotoxicity [41].

*Second, do other viral RDEs modulate target cell surfaces to a similar extent?* A range of viruses targets different host sialic acid derivatives as highly specific attachment receptors, and many of these also express a RDE [2]. The influenza A virus neuraminidase promotes viral fitness [6, 43], but little is known about its direct effects on host cells during infection.

The same is true for other viral RDEs. Two studies have documented loss of host sialic acids in lung tissues of influenza virus A (H1N1, H1N2, and H4N6)- and high pathogenic avian influenza virus (HPAIV) A (H5N1)-infected animals, as well as *ex vivo* HPAIV A (H5N1)-infected human lung biopsies [44, 45]. These studies support the assumption that influenza A virus, similar to ISAV, modulates target cell surfaces during infection. Moreover, evidence for destruction of the influenza virus human erythrocyte receptor was provided in haemagglutination reactions more than 70 years ago [46]. Still, we have not identified any studies that address the effect of influenza A infection on erythrocyte surfaces in infected humans or animals. While most influenza A strains rarely cause viraemia in humans, HPAIV A (H5N1) is an exception [47]. Considering the zoonotic and even pandemic potential of this virus [48], we suggest that the consequences of a widespread loss of virus-targeted sialic acids in this scenario should be addressed by future studies.

## Materials and methods

### Fish and experimental infection

Atlantic salmon (AquaGen, Trondheim, Norway) were hatched, reared, and kept at the aquaculture research station: Center for Sustainable Aquaculture (Norwegian University of Life Sciences [NMBU], Ås, Norway). Fish were maintained on a 24 hour light photoperiod in circular tanks in a temperature-controlled (14±1 °C) fresh water recirculatory aquaculture system and fed a standard salmon diet in excess, using automatic belt feeders. Blood for use in *ex vivo* experiments was collected in heparinised containers by terminal blood sampling from the caudal veins of fish anaesthetised by Tricaine methanesulfonate immersion (100-200 mg/L). Prior to the infection trial, the relevant fish group was tested by qPCR and found negative for infectious salmon anaemia virus (gills/heart/kidney), salmon pox gill virus (gills), infectious pancreatic necrosis virus, piscine rheovirus-1, piscine myocarditis virus, and salmonid alphavirus (all heart/kidney) by the diagnostic services at the Norwegian Veterinary Institute.

A total of 73 fish (median body weight 115 g) were transferred to the NMBU infection Aquarium (Ås, Norway) for use in the infection trial. The unit uses a fresh water flow-through aquaculture system at 12 °C. After acclimatisation, one group (n=47) was infected by 2-hour immersion in water containing the high-virulent Norwegian ISAV isolate NO/Glesvaer/2/90 [35] (10^3.75^ TCID_50_/mL), as previously described [49]. This protocol reliably infects all fish in a synchronised manner, but the severity of disease varies between trials [49, 50]. Infected fish were sampled 4, 6, 8, 10, 12, 14, 16, and 18 dpi (n=4-6 fish per time point). A group of non-infected fish from the same batch (n=26) served as controls and was sampled 0, 12, 14, 16, and 18 dpi (n=4-6 fish per time point). At sampling, fish were anaesthetised by immersion in benzocaine (40 mg/L), weighed and measured, and blood was collected from the caudal vein into heparinised containers. After blood sampling, the exterior of the fish was examined, fish were killed by cervical sectioning, the midline was incised, and organs were inspected and harvested. Haematocrits were measured within 90 min after sampling, by measuring the cell fraction of centrifuged blood (75 mm capillary tubes, 1200 rpm, 3 min, room temperature [RT]).

### Cells

Atlantic salmon kidney (ASK) cells [51] were maintained in L-15 medium (Lonza) supplemented with foetal bovine serum (FBS, 10%, Lonza), L-glutamine (Lonza, 4mM), and penicillin/streptomycin/amphotericin (Lonza, 1%) or gentamicin (50 μg/mL). The cells were cultured in closed cap tissue culture flasks at 20°C and were split 1:2 every 14 days.

Erythrocytes for *ex vivo* experiments were isolated from heparinised blood from non-infected Atlantic salmon by density gradient centrifugation. Briefly, 0.25 mL blood was diluted in 5 mL phosphate-buffered isotonic saline (PBS), layered on top of 7.5 mL 51% Percoll Plus (Sigma-Aldrich), and centrifuged without breaks (500x *g*, 20 min, 10 °C). The erythrocyte pellets were extensively washed in PBS and resuspended 2 × 10^7^ cells/mL in the same culture medium as the ASK cells. Erythrocyte suspensions were maintained in 6-well cell culture plates on a microplate shaker (IKA-Werke, 150-200 rpm, 15 °C) for up to 10 days. ISAV exposure was performed by resuspending erythrocytes in medium containing ISAV, plasma from infected fish, or recombinant HE (serial dilutions), followed by incubation in 24- or 48-well cell culture plates on a microplate shaker. Where relevant, ISAV or plasma from infected fish was pre-incubated with a mouse IgG_1_ targeting ISAV HE (clone 9G1F10A, generated by Knut Falk in the late 1990s) or mouse IgG_2a_ targeting influenza virus NP (clone D67J, Thermo Fisher Scientific) (both: 5 μg/mL, 30 min, RT).

### Virus and titering

ISAV was propagated in ASK cells at 15 °C, and supernatants were harvested when cytopathic effects were close to complete, 14-28 dpi. Infective titres were determined by inoculating serial dilutions of supernatants or blood (10 μL heparinised blood was added to 500 μL L-15 medium and kept at −80 °C until titration) in quadruplicate wells of ASK cells cultured in 96-well microtiter plates. Acetone-fixed cells were incubated with IgG_1_ against the ISAV nucleoprotein (P10, Aquatic Diagnostics Ltd, 0.4 μg/mL, 60 min, RT), washed (PBS × 3), and incubated with Alexa488-labelled goat anti-mouse IgG (A11001, Thermo Fisher Scientific, 5 μg/mL, 45 min, RT), and titres were calculated by the modified Kärber method, as previously described [17].

### RNA extraction and qPCR

20 μL heparinised blood was added to 400 μL MagNA Pure LC RNA Isolation Tissue Lysis buffer (Roche). Head kidney pieces (3 mm × 3 mm) were collected in RNA later (Thermo Fisher Scientific), transferred to 500 μL MagNA Pure LC RNA Isolation Tissue Lysis buffer, and homogenised with 3-5 mm steel beads in a TissueLyser II (Qiagen, 24 Hz, 2 × 3 min). 350 μL lysed blood or 200 μL lysed head kidney were transferred to a Magna Pure 96 Processing Cartridge (Roche), and the total volume of head kidney samples was adjusted to 350 μL by adding MagNA Pure LC RNA Isolation Tissue Lysis buffer. RNA was extracted on a MagNA Pure 96 instrument (Roche) with the MagNA Pure 96 Cellular RNA Large Volume Kit (Roche), using the RNA tissue FF standard cellular RNA protocol with an elution volume of 50 μL per sample. RNA yield and purity was determined by a Multiskan Sky spectrophotometer (Thermo Fisher Scientific). The QuantiTect reverse transcription kit (Qiagen) was used for cDNA synthesis. Real time PCR was carried out with the TaqMan fast Advanced kit (Applied Biosystems) using the following protocol: TaqMan Fast advanced master mix (1x), TaqMan assay probe (0.2 μM), forward and reverse primers (0.5 μM), and cDNA template (0.5 ng/μl) final concentrations. Thermocycling was performed on a CFX384 Bio-Rad) and CFX-manager (software version 3.1, Bio-rad) under the following conditions: 2 min at 50 ▫, 20 seconds at 95 ▫, 40 cycles of 3 seconds each at 95 ▫, and 30 seconds at 60 ▫.

### Flow cytometry

100 μL heparinised blood from each fish in the infection trial was collected in PBS, washed, and fixed (3% paraformaldehyde, 10 min, RT) before staining (RT). Erythrocytes from *ex vivo* experiments were stained live (4 °C). Cells were labelled with Alexa488-labelled wheat germ agglutinin (1 μg/mL, Thermo Fisher Scientific, 60 min), FITC-labelled sambucus nigra lectin (10 μg/mL, Thermo Fisher Scientific, 60 min), or mouse IgG_1_ specific to the ISAV hemagglutinin esterase (clone 3H6F8 [52], hybridoma supernatants diluted 1:10 for fixed and 1:100 for live cells, 60 min), washed (PBS × 3), and (for detection of ISAV only) incubated with Alexa488-labelled goat anti-mouse IgG (A11001, Thermo Fisher Scientific, 5 μg/mL, 45 min). Signal was detected by a NovoCyte flow cytometer (Agilent).

### Immunostaining of blood smears

Blood smears were made and fixed (80% acetone, 15 min, RT) within 120 min of sampling, dried, and stored at −80 °C until staining. Thawed and dried smears were incubated with 1x clear milk block (Thermo Fisher Scientific, 30 min, RT), mouse IgG_1_ specific to the ISAV HE (clone 3H6F8 [52], hybridoma supernatant diluted 1:100, 60 min, RT), washed (PBS × 3), and incubated with Alexa488-labelled goat anti-mouse IgG (A11001, Thermo Fisher Scientific, 5 μg/mL) together with Alexa594-labelled phalloidin (Thermo Fisher Scientific, 2 units/mL, 45 min, RT). Nuclei were counterstained by Hoechst 33342 (Thermo Fisher Scientific, 2 μg/mL, 5 min, RT), and slides were mounted in ProLong Gold antifade mountant (Thermo Fisher Scientific). Wide-field microscopy was performed by a Nikon Ti-2e inverted microscope, using a Plan Apo lambda DIC N2 63x oil objective (NA 1.4). The percentage of ISAV-positive cells in individual fish was quantified in ImageJ (version 1.53c) [53]: Total cell numbers were determined using the *Analyze particles* function on thresholded images generated from the Hoechst channel; next, the number of ISAV-positive cells were counted manually in the channel containing signal from the HE-staining, assessing a minimum of 99 cells per fish.

### Histology

An organ panel including heart, spleen, and head kidney was collected from each fish, fixed in 10% formalin for at least 24 hours, dehydrated, and embedded in paraffin. Thin tissue sections were heat treated (60-70 °C, 20 min), deparaffinised, and either stained with haematoxylin & eosin for histological evaluation or subjected to a virus binding assay, as described below.

### Preparation and blotting of membrane-enriched lysates

Cell pellets were prepared from full blood or density-purified erythrocytes and stored at −80 °C before preparation of membrane-enriched fractions as previously described [24]. Briefly, 100 μL cell pellets were lysed by 1:10 dilution in ice cold water with 1% protease inhibitor (P8340, Sigma-Aldrich, 10 min, on ice). The cells were homogenised with a tight-fitting dounce homogenisator (20 strokes), 1000 μL buffer A (75 mM Tris pH 7.5, 12.5 mM MgCl, 15 mM EDTA) was added, and the homogenisation was repeated. To remove nuclei and organelles, the suspension was centrifuged (5000× *g*, 5 min), and the supernatant was collected on ice. The homogenisation procedure was repeated three times with the cell pellet in buffer A diluted 1:1 with water, pooled supernatant fractions were centrifuged (40,000× *g*, 30 min), the membrane-enriched pellets were resuspended in 50 μL buffer B (20 mM Tris, 2 mM EDTA, pH 7.5), and samples were stored at −80 °C. After thawing, 10 μL sample, 3.88 μL NuPage LDS sample buffer, and 1.55 μL NuPage sample reducing agent (both from Thermo Fisher Scientific) was mixed and homogenised by centrifugation in QiaShredder columns (Qiagen), heat-treated (10 min, 70 °C), and loaded to gels (8 μL/well) for SDS-PAGE and Western blotting (NuPage Novex system, Thermo Fisher Scientific) to 0.45 μm nitrocellulose membranes (BioRad laboratories).

### Production of antigen for virus binding assays

Antigens for virus histochemistry and virus binding assays were prepared by collecting membrane fractions of infected ASK cells, as previously described [21]. Briefly, cells in 75 cm^2^ tissue culture flasks were harvested when cytopathic effects were evident but most cells remained attached, typically 3-7 dpi. Cells were washed and scraped on ice, and cell pellets containing cellular-expressed viral membrane glycoproteins were washed, resuspended in 0.5 mL PBS, and subjected to three rounds of freeze-thawing. Hemagglutination titres were determined by incubating serial dilutions of antigen preparations with 1% Atlantic salmon erythrocyte suspensions in 96-well V-bottom microtiter plates.

### Virus binding assays

To map virus-binding sites in Atlantic salmon tissues and membrane-enriched cell fractions, we used virus antigen preparations as primary probes as previously described [21, 23, 24].

Deparaffinised formalin-fixed tissue sections were incubated with 100 μL ISAV antigen (512 HAU/mL, 60 min, RT), washed (PBS × 3), quenched with peroxidase block (5 min, RT), treated with blocking buffer (Background sniper, Biocare Medical, 30 min, RT), incubated with mouse IgG_1_ specific to ISAV HE (clone 3H6F8 [52], hybridoma supernatants diluted 1:100, 60 min, RT), washed (PBS × 3), and signal was visualised by the MACH2 HRP polymer-DAB (Biocare Medical) system, following manufacturer’s instructions. Signal was analysed as described in the section *Quantification of virus histochemistry signal*.

Blotted membrane-enriched cell lysates (prepared as described above) were stained for total protein using Revert700 Total Protein Stain as recommended by the manufacturer (Licor Biosciences), treated with dry milk (5% in Tris-buffered saline [TBS], 60 min, RT), washed (TBS 0.1% Tween [TBST] × 2), incubated with ISAV antigen (256 HAU/mL diluted in TBS, 60 min, RT), washed (TBST × 3), incubated with mouse IgG_1_ specific to ISAV HE (clone 3H6F8 [52], hybridoma supernatants diluted 1:250 in TBST with 5% dry milk, 60 min, RT), and washed (TBST × 3). Bound primary antibody was detected either by incubation with IRDye800-labelled goat anti-mouse IgG (Li-Cor Biosciences, 1:10,000 in TBST with 5% dry milk, 60 min, RT), or by incubation with HRP-conjugated horse anti-mouse IgG (Cell Signaling, 1:1000, 60 min, RT), wash (TBST × 3), and incubation with Super Signal West pico substrate (Thermo Fisher scientific). Chemiluminescent or fluorescent signal was visualised by an Azure imager C500 (Azure Biosystems). For quantification, fluorescent signal was normalised to total protein signal, using Azure spot software (Azure Biosystems, version 2.2.167).

### Quantification of virus histochemistry signal

Images of tissue sections subjected to virus binding assays were digitized using a Hamamatsu Nanozoomer digital slide scanner with 40x objective (114932 pixels per inch, 221 nm/pixel, JPEG compression at 80%). The virus histochemistry signal was analysed using Visiopharm, v2022.2. For region of interest (=organ) detection and area measurement, a generic analysis protocol package using artificial intelligence (DeepLab V3) trained in our lab on collections of slides containing Atlantic salmon liver, heart, kidney, spleen and adipose tissue, was retrained on the full set of stained sections from the present study. Signal detection was performed at 20x magnification: The image pixel value was inverted, multiplied by 5, and combined with their red-green contrast values using a maximum pixel value filter. Pixels with resulting values above a threshold of 50 were considered positive. The total area covered by positive pixels per organ was divided by organ area to yield the fraction of positive signal relative to area of the organ in the section.

### Generation of recombinant ISAV HE

Codon optimised sequences of the open reading frame encoding ISAV HE, corresponding to isolate NO/Finnmark/NVI-70-1250/2020 (Genbank accession UGL76651.1), as well as a variant where serine 32 was mutated to alanine to abolish catalytic activity (S32A), was synthesised and inserted in the pcDNA3.1 (+) vector commercially and delivered transfection-ready (GeneCust). Monolayers of CHSE-214 cells were cultured until 90-100% confluent and detached by Trypsin EDTA (Lonza). 4 transfection reactions, each with 10^6^ cells and 10 μg plasmid DNA, were performed, using the Neon 100 μL Transfection System (Invitrogen, Waltham, MA, USA, three pulses at 1600 V and 10 ms width). Non-transfected cells were used as controls. The transfected cells were pooled and incubated 24 hours in antibiotic free medium, then another 24 hours in culture medium. Membrane fractions were collected as described for infected cells and previously [21].

### Statistics

Statistics were performed in Graph Pad Prism 8 for Windows 64-bits, v.8.4.3.

### Ethical considerations

Protocols for harvesting material from healthy fish and experimental infection and their implementation were approved prior to the studies by the Norwegian Food Safety Authority (FOTS24383, FOTS28403). All facilities were operated in compliance with Organisation for Economic Co-operation and Development principles of Good Laboratory Practice and Guidelines to Good Manufacturing Practice issued by the European Commission.

## Acknowledgements

The authors would like to thank: Eirill Aager-Wick and Hetron Mweemba Munang’andu (the NMBU infection aquarium for salmonids, Ås, Norway) for administrative and technical support for the infection trial; Ricardo Tavares Benicio and Bjørn-Reidar Hansen (the aquaculture research station, NMBU, Ås, Norway) for providing fish and technical assistance; Torfinn Moldal (Norwegian Veterinary Institute) for organising the pre-trial screening for infectious agents; Marit Måsøy Amundsen, Randi Faller, Randi Terland, Britt Saure, Lone Engerdahl, and Ingebjørg Modahl (Norwegian Veterinary Institute) for technical assistance; and Maria Aamelfot (Norwegian Institute for Public Health) for fruitful discussions. The authors would also like to acknowledge two research projects funded by the Research Council of Norway (grant 302551) and strategic base-funding at the Norwegian Veterinary institute (BIODIRECT: Biomarkers and bioassays for future research and diagnostics) for methodological support and refinement.

## Supplemental materials

**S1 Table: Numericals and statistics used to generate the figures**

**S1 Fig, supplemental to Fig 1:** (A-B) Manual scoring and (C) example micrograph of erythrophagocytosis in (A) head kidney and (B-C) spleen in haematoxylin and eosin-stained formalin-fixed, paraffin-embedded tissue sections of individual fish. Arrows point to examples of macrophages that have ingested eosinophilic debris, considered evidence of haemophagocytosis. (D) Viral RNA in head kidney was measured by qPCR targeting ISAV segment 8. Data points show values in individual fish, the line connects median values.

**S2 Fig, supplemental to Fig 5:** (A) ISAV was pre-incubated with monoclonal antibodies as indicated (5 μg/mL, 30 min, RT), before inoculation with density-purified erythrocytes isolated from non-infected fish (10^6^ TCID_50_ per 2 ×10^7^ cells, 60 min) and quantification of HE by flow cytometry. Data points show measurements in cells from individual fish. **p<0.01, Kruskal-Wallis with Dunn’s multiple comparisons test (S1 Table). (B) Haemagglutination inhibition assay testing the ability of 9G1F10A to inhibit ISAV-induced agglutination (4 HAU ISAV antigen and 10^6^ erythrocytes per well). The boxed area indicates the concentration range of the antibody that completely inhibited agglutination.

