## Supplementary figures and images for "The infectious salmon anaemia virus esterase prunes erythrocyte surfaces in infected Atlantic salmon and exposes terminal sialic acids to lectin recognition"

### S1 Fig

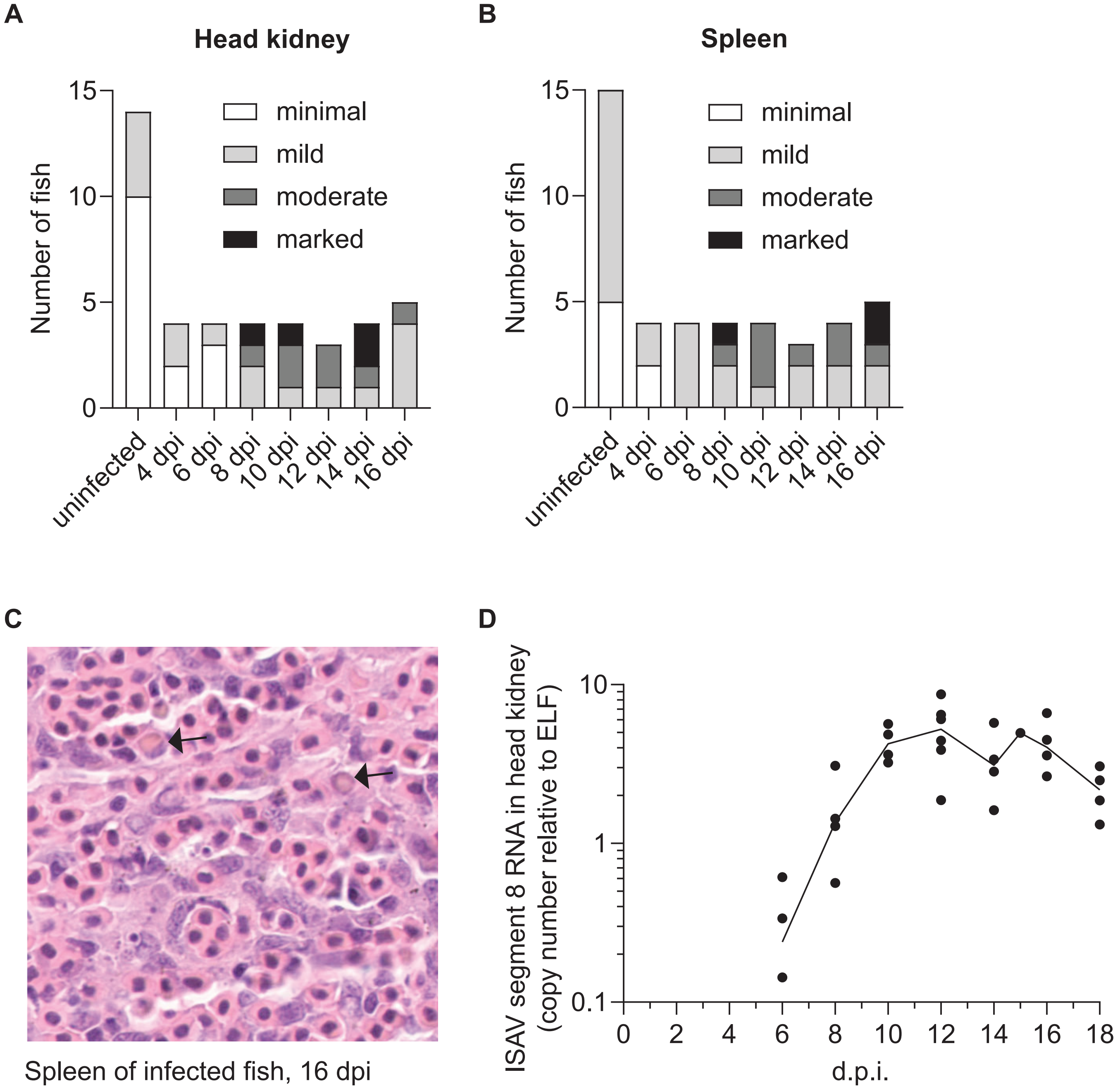

### S2 Fig

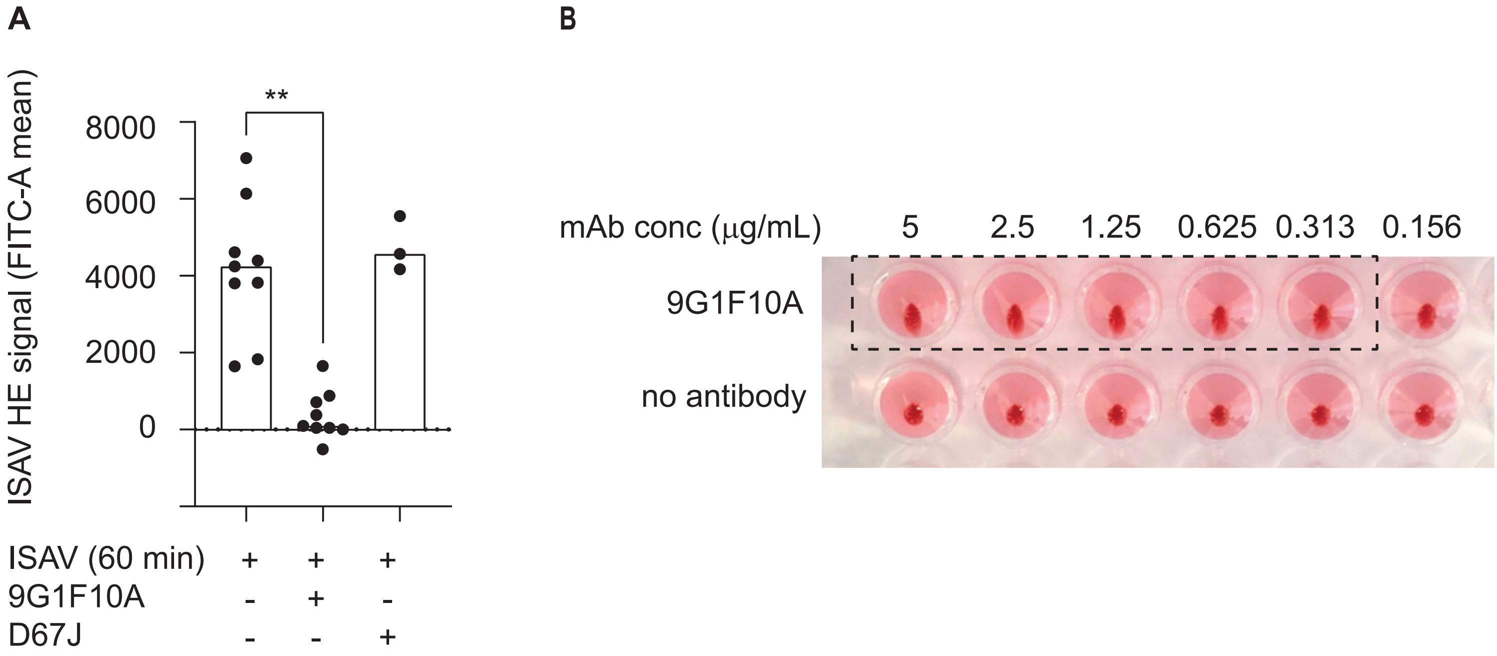
